# Response to “Commentary on ‘Limitations of GCTA as a solution to the missing heritability problem”

**DOI:** 10.1101/039594

**Authors:** Siddharth Krishna Kumar, Marcus W. Feldman, David H. Rehkopf, Shripad Tuljapurkar

## Abstract

In a recent manuscript [18], Yang and colleagues criticized our paper, ‘Limitations of GCTA as a solution to the missing heritability problem’ [9]. Henceforth we refer to our original paper as SK and the critique as YC.

Here we show that their main claims are statistically invalid. We begin with an overview and the mathematical details follow in subsequent sections.

## 1 Overview

YC begin their critique with a qualitative description of how they believe the genotype ought to influence the phenotype. That discussion motivates the statistical model they use to estimate heritability. In our analysis (SK), we do not contest their motivation for constructing the model (e.g., their view that some tagged SNPs are in strong LD with the causal SNPs). We simply analyze their mathematical model, and show that it produces unreliable estimates. We first summarize our arguments, highlighting material omitted from SK because of space restrictions.

### The GCTA model

A GWAS is summarized in a ‘genotype’ matrix, **Z** for *N* individuals and *P* SNPs, and we want to know how this genotypic information influnces some particular phenotype. GCTA models the phenotype vector of the *N* individuals, y as

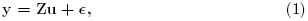

where the vector, **u** ~ 𝒩(0, *σ*^2^**I**) and ϵ ~ 𝒩(0, *α*^2^**I**). This equation can be rewritten as

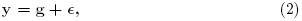

where **g** ~ 𝒩(0, *σ*^2^**A**), and **A** = (1/*P*)**ZZ**^**T**^ is the Genetic Relatedness Matrix (GRM). In reality, we only have an estimate of the genetic structure (for many reasons, e.g., the population of people is a sample from some larger population; the same holds for the SNPs sampled; we may not be able to accurately call the state of all SNPs for all individuals; some SNPs make no contribution or unequal contributions to the phenoype) described in (1). Therefore, we assume in SK that **Z** (correspondingly, **A**) differs from a true genotype matrix (GRM), **Z**_1_ (correspondingly, **A**_1_) by some estimation error, **E**(correspondingly **F**) so that

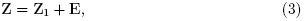

and

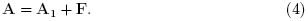

Equations (1)-(4) provide a precise mathematical description of GCTA; the assumptions made by GCTA follow from the mathematical restrictions enforced by the model. The key assumptions of GCTA and our (SK) deductions from them are:

- GCTA assumes that the covariance matrix of u is diagonal, hence every SNP used in the GCTA model (irrespective of whether it is in strong LD with a causal SNP) necessarily makes a random contribution to the phenotype. Non-causal SNPs would make zero contributions to the phenotype, which would require that the corresponding diagonal entries of the covariance matrix of **u** be zero; GCTA does not set any diagonal entry to 0 in its formulation.
- Equation (1) is not stated as conditional on *N* or *P* and therefore *σ^2^* is also not conditional on *N* or *P*. YC state that *σ^2^* must be interpreted as the ‘variance of SNP effects when it is fitted jointly with all other SNPs’. However, in the model as stated, *σ^2^* is the variance of a random contribution made by each SNP, irrespective of which other SNPs are used in the analysis. We discuss this point in greater detail below in section 2.
- Since the model formulation involves no specification of **Z**|**Z**_1_, the distinction between causal and non-causal SNPs is irrelevant to the application of GCTA.
- Since the off diagonal entries in the covariance matrix of u are assumed to be 0, GCTA necessarily assumes that the random contribution made by each SNP is not correlated with the random contributions made by any other SNPs used in the analysis. In SK, we interpreted this assumption to mean that the SNPs are in linkage equilibrium with each other, regardless of their linkage status with the (unknown) causal SNPs.

### Our analysis of GCTA

In SK, we used population genetics, theory of random matrices and numerical analysis to show that the heritability estimates produced by GCTA will be unreliable.

### The data used in SK

In SK we used the Framingham (FHS) Share dataset containing information on *N* = 2, 698 ‘unrelated’ individuals and *P* = 49, 214 SNPs. For our analysis, we used pedigree information in the FHS to retain only those individuals who did not share the same parents (i.e., the relatedness is reduced to at most first cousins). We also filtered our data to remove missing entries using an analysis similar to what is done in table 2 of [11]. The phenotype data we used in our paper is taken from study accession phs000007.v23.p8.

We argue that (a) the singular values of **Z** are skewed i.e., the largest singular value of **Z** (or eigenvalue of **A**) is much larger than the smallest singular value of **Z**, and that (b) the smallest singular values of **Z** are close to 0. This is not a novel observation; the skews in the singular values for the Framingham dataset have previously been reported in Figure 4 of Hoffman [7] where the eigenvalues of the GRM, **A** were analyzed for different datasets.(That figure is reproduced here as Fig. 1, with permission). In Fig. 1 the eigenvalues fall sharply to an intermediate value, but eventually fall to values close to 0 (recall: eigenvalues are (1/*P*) times squared singular values). Importantly, the ratio of the largest to the smallest eigenvalue value of the sample GRM **A** (or the ratio of the largest to smallest singular value of **Z**) is large.

**Figure 1:**
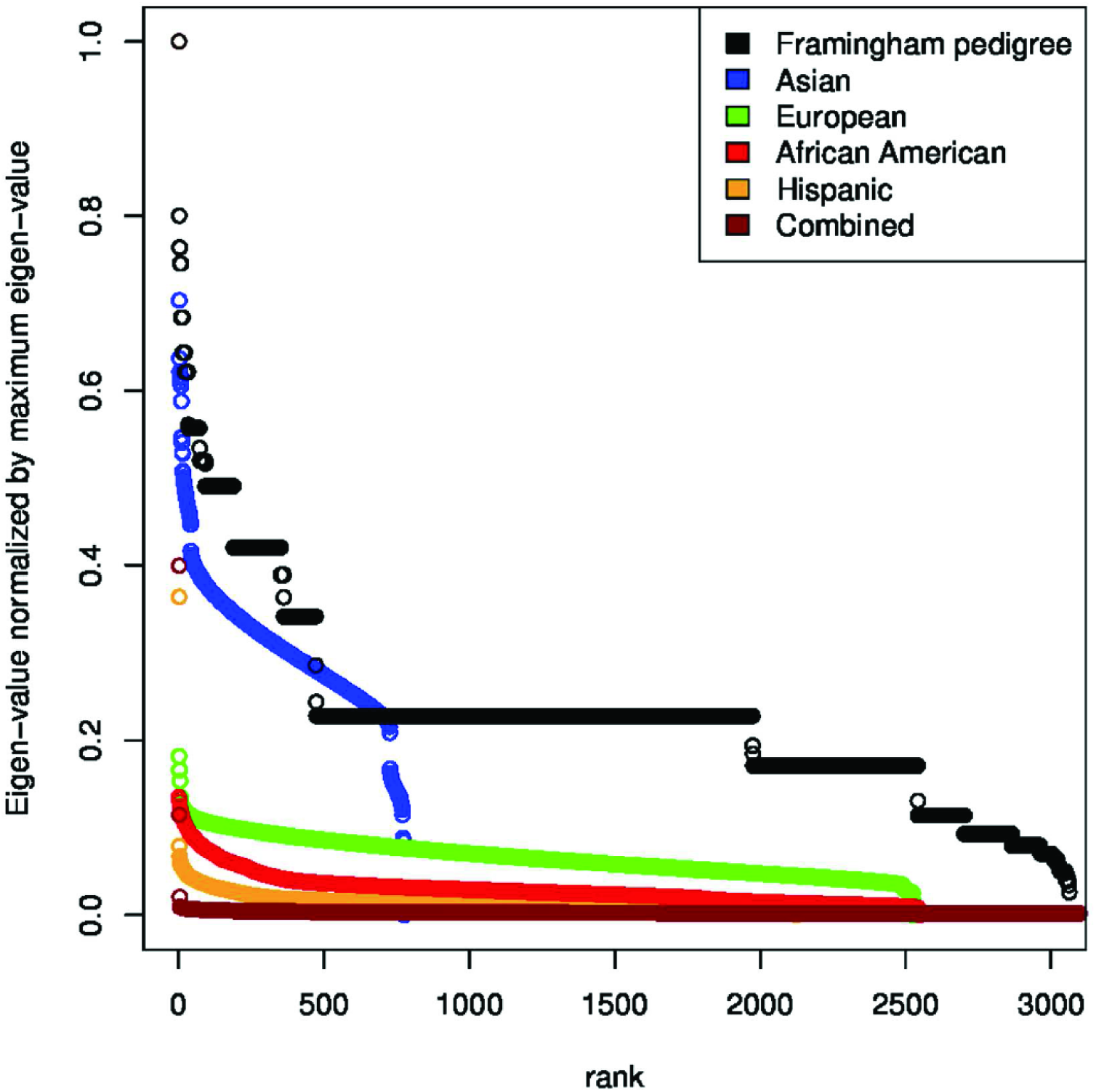
Skews in singular values for different populations. This is Fig. 4 in [7].

**Figure 2:**
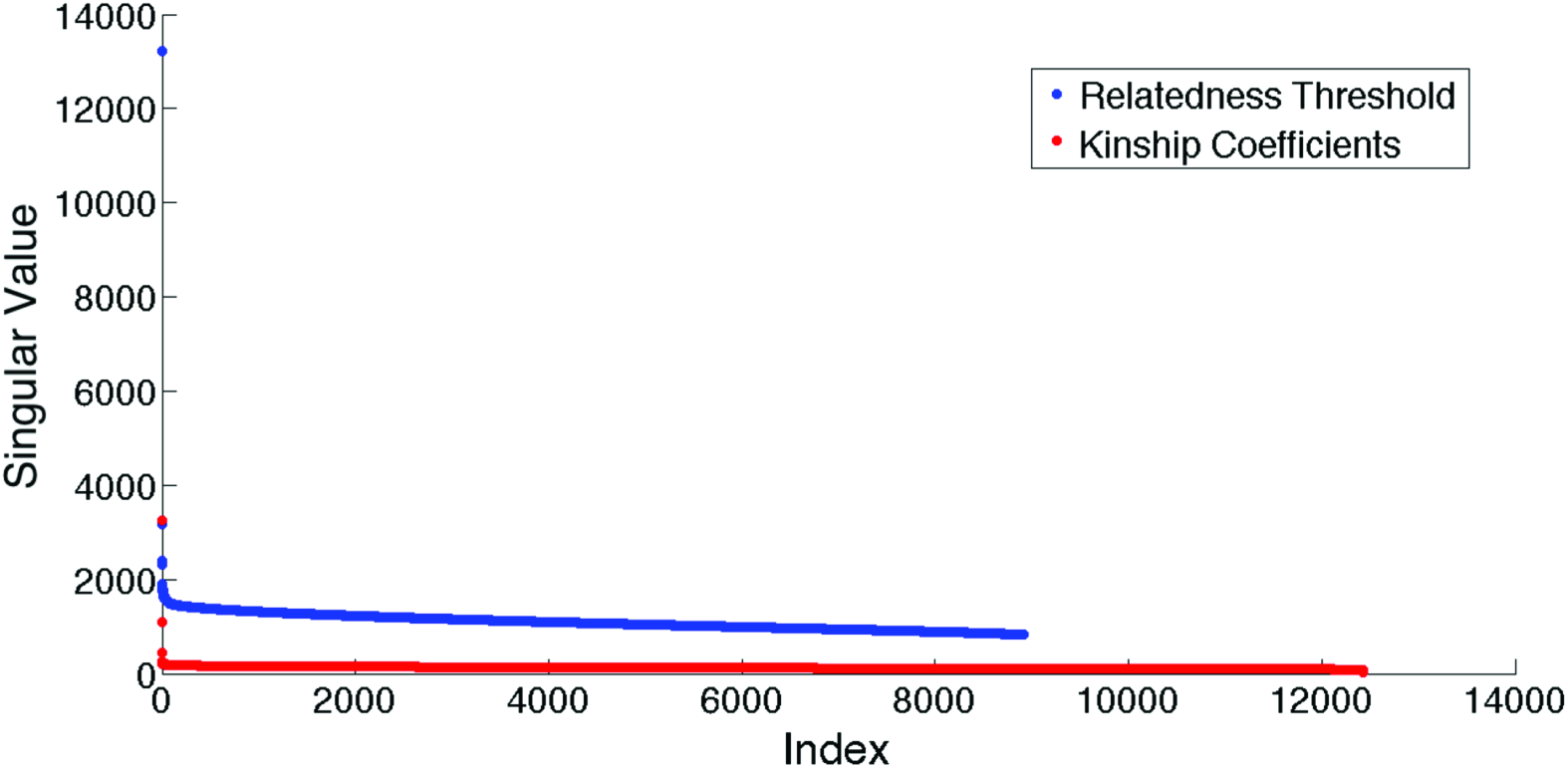
Skews in Singular values when using samples filtered using kinship coefficients (in red) and using relatedness threshold (in blue). Notice that there is a ‘gap’ between the red and blue dots being caused because of cryptic relatedness.

**Figure 3:**
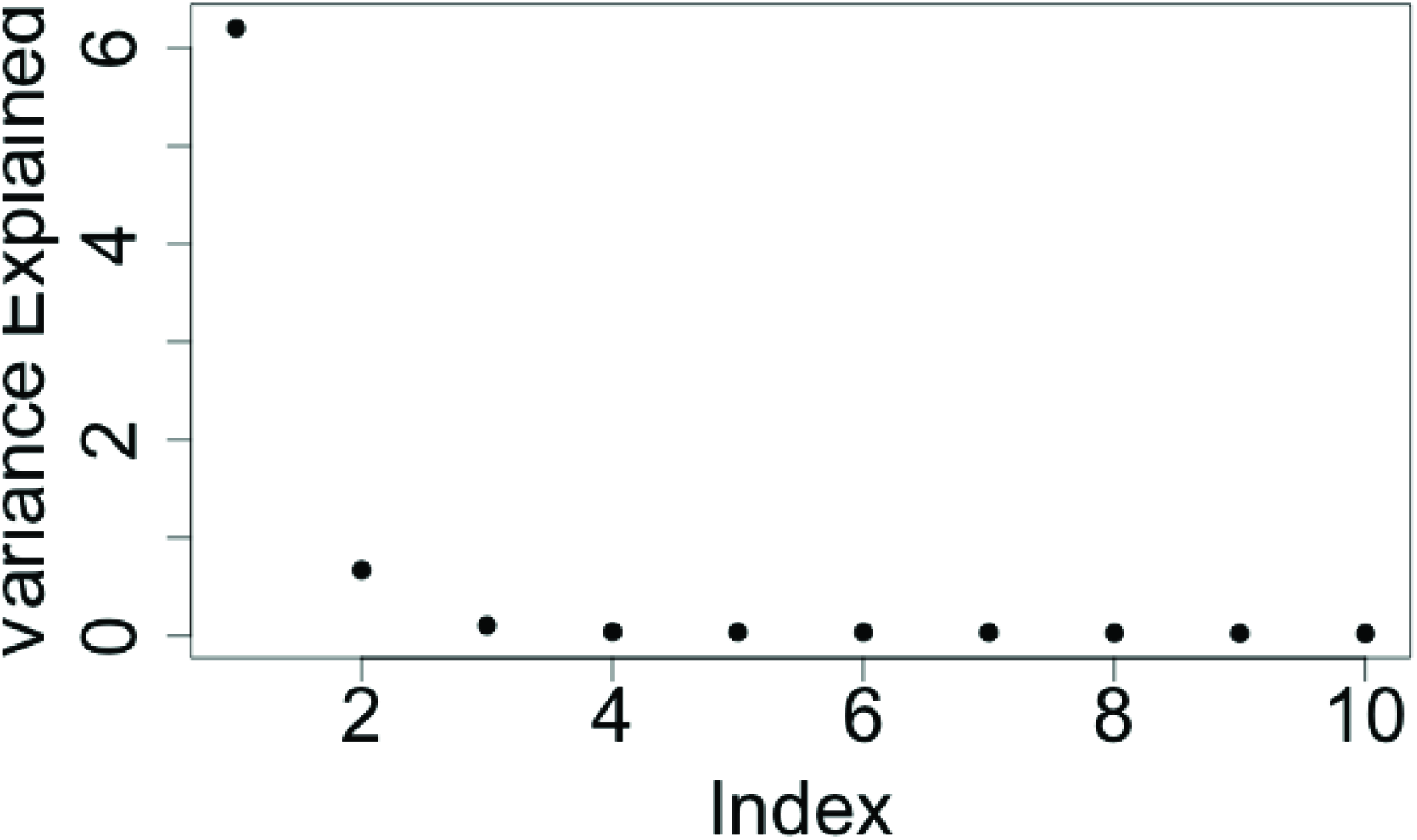
First ten eigenvalues of the GRM for the HRS dataset.

**Figure 4:**
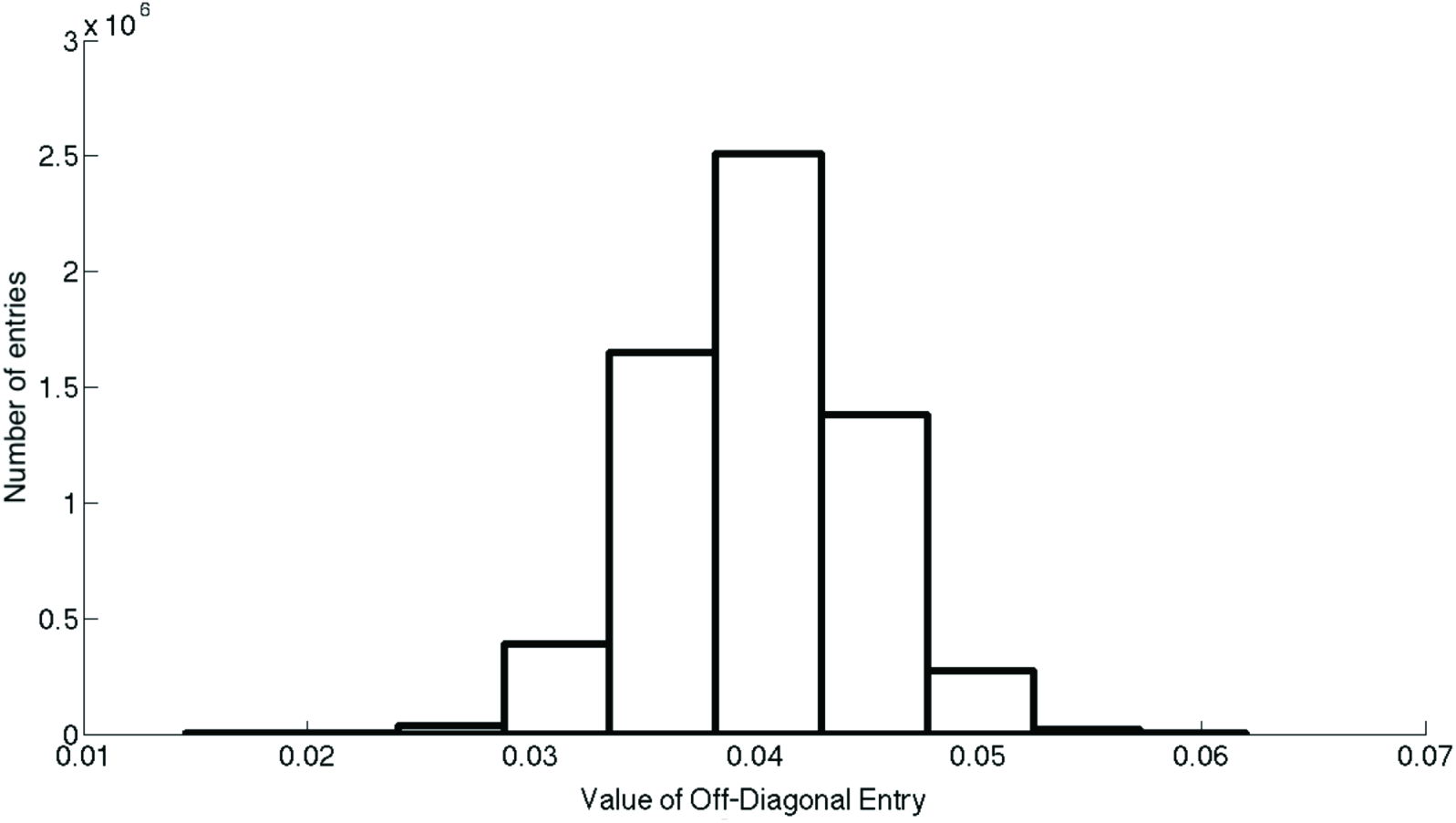
Histogram of off-diagonal entries of sample covariance matrix when the ‘true’ covariance matrix has all off-diagonal entries = 0.04. Note that there are some entries with errors greater than 50%

The skew we report for the singular values from FHS is to be expected given the theoretical work in [12]. Similar skews have been reported from analysis of other datasets (see the eigenvalue distributions for the Asian, European, African American and Hispanic populations in Fig. 1). The skew observed in the Framingham dataset is not a special case: to demonstrate, we plot in Fig. 2 the skew in the singular values (marked with red dots) derived from the Health and Retirement Study (HRS) dataset (quality control instructions are provided in http://hrsonline.isr.umich.edu/sitedocs/genetics/HRS_QC_REPORT_MAR2012.pdf). For clarity, in Fig. 3 we also provide the first 10 eigenvalues of the GRM (matching exactly Figure 16 of the link above). The ratio of the largest to smallest eigenvalue of the GRM in the HRS dataset is 6.7×10^18^, which is the same order of magnitude reported for FHS in our paper (SK).

The smallest eigenvalue in the GRM for the HRS dataset is so small that it falls below machine accuracy in our computation, and is therefore reported as −4.3 × 10^−17^. Since the GRM is positive semi-definite, we assume the smallest value is 4.3 × 10^−17^, although the true value might be a smaller positive number. It is important to note that the smallest eigenvalue of the GRM is near-zero, but not precisely zero. We know from (a) Theorem 3 in [12] that the smallest eigenvalue of the true GRM is precisely 0, and (b) from standard results in perturbation theory [15] that the smallest eigenvalue of the true GRM and sample GRM will not be exactly the same.

YC analyze the FHS data, and report that when they chose ‘unrelated’ individuals using the filtering techniques in [17] (which they call relatedness thresholding), they do not find large skews in the singular values. Furthermore, they do not find any singular values close to 0 and accordingly conclude that we have made a mistake in our analysis. We show in section 4 below that YC do not observe the skew because the filtering technique suggested in [17] is flawed.

### A description of our analysis

In SK, we analyzed the likelihood function used in the GREML and showed that it produces unstable estimates even when the assumptions in GCTA are satisfied exactly. The instabilities arise because the singular values of **Z** are packed close to one another. Furthermore, we showed that when there is population stratification (as is to be expected in most real datasets), the instabilities can be severe because the ratio of the largest to smallest eigenvalue of the GRM will be large.

YC do not address the problems we raise regarding the close packing of the singular values, but do address our claims of large skews in the singular values of **Z**. They claim that even when present, such skews will not cause instabilities in the estimates. In Section 5 below, we provide a precise mathematical description of why large skews in the singular values of **Z** make the GREML estimates sensitive to even small sampling errors; we use published results to show that the sampling errors in the GRM are large and therefore, the estimates produced by GCTA are unreliable.

In SK, we showed that GCTA overfits the data because it estimates *O*(*NP*) parameters from a dataset containing *NP* entries. In such a setting overfitting is guaranteed, and is a direct consequence of the bias-variance tradeoff, In their critique YC simply state that GCTA is not overfitting; they try to show that GCTA is not overfitting by comparing it to seemingly similar statistical models such as linear regression and mixed linear models. We show that in Section 3 below that, in contrast to these methods, GCTA omits either (a) important conditioning arguments or (b) procedures to prevent against overfitting.

### Our simulation experiments

SK demonstrated the instabilities in the GCTA estimates by producing a contradiction: we constructed subsets of SNPs from an overall sample, and showed that several of the *σ^2^* estimates for these subsets *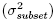* lie outside
the 99.5% confidence intervals predicted from the estimates and standard errors of *σ^2^* for the overall sample *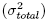*. YC claim that our analysis is flawed because 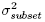 is not, in their view, the same parameter as 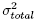. However, as we explain below, the equality of 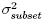 is and 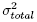 is precisely what we do expect. We also used published results to demonstrate that GCTA’s estimates are sensitive to the sample of people used in the study, and to population stratification.

## 2 What does GCTA estimate?

Suppose the *P* SNPs which satisfy GCTA’s assumption exactly are known (i.e., the true genotype matrix **Z**_1_ is known). Then the variance in the phe-notype can be be estimated precisely as

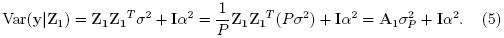

where **A**_1_ is the true GRM (corresponding to **G** in [17]). Thus GCTA actually assumes that

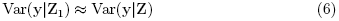

for all **Z**. This implies that

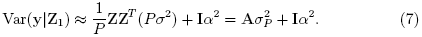

Comparing (5) and (7) shows that GCTA requires **A**_1_ ≈ **A** irrespective of which *P* SNPs are used in the analysis. This requirement is satisfied only when all SNPs in the data set make independent and identically distributed contributions to the phenotype. This is the assumption we state in SK.

It is important to note that GCTA does not assume that the observed **A** is an *estimate* of the true **A**_1_. The latter assumption would require treating **A**|**A**_1_ as a random variable, which would invalidate the variance calculation in (7).

Put another way, the SNPs included in any GWAS are determined by the technology and not by prior information about the causal or non-causal relationship between these SNPs and the phenotype under study. Thus we should expect to get the same GCTA estimates of variance from a data set on *N* people obtained by using say five chips each with 500,000 distinct and non-overlapping SNPs, or by using 1 chip with all 2.5 Million SNPs. Thus we do expect that 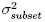 and 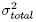 describe the same parameter, and since the number of individuals in both studies is the same, we expect the two parameters to produce near-identical confidence intervals.

Furthermore, since GCTA’s model assumes that the variance contribution of each SNP is the same, we expect the total variance contribution of all the SNPs used in the analysis to be directly proportional to the number of SNPs. We demonstrated a violation of this expectation in the section titled ‘Saturation of heritability estimates’ in SK.

The total genome-wide variance cannot be compared across different studies because the number of SNPs used in different studies is different; these comparisons often produce misleading results. To illustrate this point, we use results from page 520 in [19] where a heritability estimate of 0.448(0.029) was obtained for height using data on 565,040 SNPs for 11,586 unrelated individuals and it was concluded that this estimate is in line with the heritability estimate of 0.445(0.083) obtained using GCTA on an Australian cohort (3,925 unrelated individuals genotyped at 294,831 SNPs). The *σ*^2^ estimate for the first study is 7.92e-7(5.1e-8), and the *σ*^2^ estimate for the Australian cohort is 1.5e-6 (2.81e-7). These two *σ*^2^ estimates are 15 standard errors apart ((1.5e-6 - 7.92e-7)/5.1e-8). These two estimates also have non-overlapping 99.5% confidence intervals of *σ*^2^ and are certainly dissimilar.

SK reports large differences between 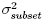 and 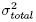. Similar large deviations have been reported [11] in studies conducted by some of YC’s authors. In table 6 of [11], for a missing genotype threshold of 200 (which means that the analysis includes only those SNPs for which there are at most 200 missing entries among the people being studied), for all SNPs without chromosome 6, GCTA produces an estimate of 0.23(0.07) for *h*^2^ on 297,028 SNPs, which corresponds to a *σ*^2^ estimate of 7.75 e-7 (2.3e-7). When only the 21,016 SNPs on chromosome 6 are used, GCTA produces an estimate estimate of 0.33(0.02) for *h*^2^; this corresponds to a *σ*^2^ estimate of 1.58e-5(9e-7). The two *σ*^2^ estimates are (1.58e-5 - 7.75e-7)/2.3e-7 = 65 standard errors apart, and produce non-overlapping 99.5% confidence intervals for *σ*^2^

### 2.1 Sensitivity to people used in the study

YC claim that GCTA does not overfit the data because the parameter being fitted in their model is random, and not fixed. However, overfitting has nothing to do with the parameter being fixed or random; it is a reflection of the bias-variance tradeoff. When the data are fit using the entire training sample, the estimation error is always going to be optimistic (see section 7.4 in [5]) and therefore, the confidence intervals produced by the model understate the uncertainty in the parameter being estimated. Cross-validation is essential, no matter what the nature of the parameter being estimated.

Overfitting produces counter-intuitive results. For example in supple-mentary table S2 in [16], the heritability of neuroticism was computed using GCTA for a sample of 5,016 unrelated men, a sample of 6,945 unrelated women, and the combined sample of 11,961 individuals; all three computations used the same 849,801 SNPs. The heritability estimate for just the women is 0.057 (0.05) and for just the men is 0.161 (0.07). The *h*^2^ estimate for the men is (0.161-0.057)/0.05 = 2.08 standard errors away from the *h*^2^ estimate of the women (since the number of SNPs used in all three studies is the same, here we can directly compare the *h*^2^ estimates instead of *σ*^2^ estimates). Furthermore, GCTA reports that the *h*^2^ estimate for the overall population is 0.056, which is smaller than the *h*^2^ estimates for both men and women.

A more extreme example is found in supplementary table S5 of [4], where estimates of the heritability of Bi-Polar Disorder (BPD) were compared for three sub-cohorts using information on 995,971 SNPs. For Subcohort 2 (2,540 cases and 2,058 controls), an *h*^2^ estimate of 0.44 (0.07) was reported, and for Subcohort 3 (1,699 cases and 2,915 controls), the *h*^2^ estimate was 0.73(0.06). These estimates two estimates are (0.73-0.44/0.07)≈4 standard errors apart and produce non-overlapping 99.5% confidence intervals for *σ*^2^.

### 2.2 Using Principal components as fixed effects

Our criticism of GCTA stems from instabilities in the eigenvalues/vectors of the GRM. Since fixed effects terms (like principal components) do not change the spectral properties of the GRM, they will not solve the problems addressed in our paper.

## 3 GCTA and similar statistical models

In their critique YC, claim that GCTA does not overfit by comparing it to other popular statistical methods. In this section, we will show that GCTA has little in common with the methods described in their critique. We show that at best, GCTA is an extreme generalization of linear regression. The standard regression model is stated as

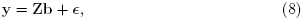

where **y** is the dependent variable, **Z** is an *N × r* data matrix describing the entries of *r* independent variables (*Z*_1_, *Z*_2_…*Z_r_*) for the *N* individuals used in the study, **b** is a vector of regression coefficients, and **ϵ** is a vector describing the errors.

There are 2 paradigms for viewing the standard linear regression model [8]. The most popular one is the ‘fixed **Z**’ paradigm according to which the regression function *E*(y|**Z**) is assumed to be linear. Here **Z** is considered ‘fixed’ and y is considered random; for the same ‘fixed’ values of **Z**, different values of y are obtained as a result of sampling errors due to **ϵ**.

The less frequently used ‘random **Z**’ paradigm assumes that the predictors and phenotype constitute an *r* + 1 dimensional random vector, with assumptions about the joint distribution providing equivalence with (8). It is well known (see pg. 112 in [8]) that although the descriptions differ as to how the data are generated in the ‘fixed **Z**’ and ‘random **Z**’ cases, the ordinary least squares estimator of b turns out to be the same in the two cases. Although the structural form of (8) seems similar to that used in GCTA, there are important differences between these two models

- In linear regression, the *r* predictors are the same in the training and test sets. However, in GCTA, the predictors are actually sets of SNPs; hence the predictor variables for different studies (that use different sets of SNPs) can be different.
- For linear regression, it has been shown that the estimate obtained in the ‘Fixed **Z**’ and ‘Random **Z**’ cases are the same. No such results are available for mixed effects models (i.e., when **b** is assumed to be sampled from a multivariate normal distribution).

### What is known for mixed effects models?

Statistical insights into mixed effects models come from the pioneering work of Henderson [6]; a survey of Henderson’s original work and several extensions are given in [14]. As stated in [14] and [13] (cited in the YC critique), these models assume that the GRM is *known* and not *estimated*. This may be reasonable in animal breeding because individuals used in the study are closely related and detailed pedigree information is commonly available. A known GRM corresponds to the case of *N* fixed and *P* → ∞ and in this limit, there are no sampling errors and therefore our analysis does not apply. Henderson’s model can be stated as

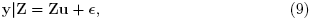

where, as before, **u** ~ 𝒩(0, *σ*^2^**I**) and e ~ 𝒩(0, *α*^2^**I**). Often, the formulation in (9) is loosely stated as

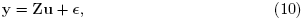

but it is made clear that **Z** is a fixed matrix. For this formulation, Henderson has shown that the estimate of *σ*^2^ will be the Best Linear Unbiased Predictor
(BLUP). Note that (9) has strong similarities with the ‘fixed **Z**’ case in linear regression - the predictors and the people used in the study are fixed.

In GCTA, **Z** is not *fixed*, but an *estimate* of the true value, **Z**_1_. Therefore, although GCTA uses a formulation that looks similar to that studied by Henderson, the equations are in fact completely different and therefore, analogies between the two models cannot be valid.

In the next section, we will show that the reason YC do not observe the skew in singular values observed in SK is that they are using the ‘relatedness thresholding’ suggested in [17] to select unrelated individuals, and this method is flawed.

## 4 Issues with Cryptic Relatedness

GCTA claims to correct for cryptic relatedness by removing all individuals whose relatedness to another is greater than some threshold (0.05 or 0.1); these thresholds are based on expectations about the true GRM, **A**_1_. However, large entries in the sample GRM, **A** do not imply that the corresponding entries in the ‘true’ GRM are large, when *N* and *P* are large. As stated in the introduction of [10], when *N* and *P* are large, the extreme values of the GRM ‘tend to take on extreme values not because this is “the truth”, but because they contain an extreme amount of error’. Therefore, the largest entries in the GRM will be most unreliable and the ‘relatedness thresholding’ done by GCTA becomes subject to a lot of error.

To expand on this issue, consider a multivariate normal distribution for a 2500-dimensional vector with mean 0 and covariance matrix **P** (of dimension 2, 500 x 2, 500, corresponding to the ‘true’ GRM) with diagonal entries equal to 1 and all off-diagonal entries equal to 0.04. We sample 50,000 random vectors from this distribution, each of dimension 2500 x 1, and compute the sample covariance matrix 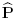.

A histogram of the off-diagonal entries of 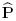 is plotted in Fig. 4. The largest off-diagonal entry in 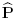 is 0.062, and the smallest is 0.0146; these extremes correspond to errors of more than 50% of the true value (which is 0.04). To apply GCTA’s threshold of 0.05 to this sample matrix, several hundred observations would have been excluded from the analysis because they would have incorrectly been deemed ‘related’. On a similar note, the small entries in the sample GRM do not mean that the subjects are ‘less related’. This example demonstrates the well known fact (see the introductions of [3],[1], [2]) that when *N* and *P* are large, no reliable inferences can be drawn from entries in the sample covariance matrix.

Since GCTA performs its relatedness filtering on the basis of entries in the sample GRM, we expect the filtering to perform poorly, and this is indeed the case.

### 4.1 The problems with Figure 1 of YC

From the genotype matrix, **Z**, we can only compute the singular values 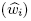 of the sample GRM, **Z**. However, we are interested in the singular values *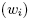* of the true GRM, **Z**_1_. For GCTA to work reliably, the error, ie., the difference between the singular values of the sample GRM 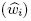, and the true GRM *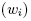* should be small. Applying Weyl’s inequality (see equation 2 in [15]), we can bound this error in eqn. (3) as

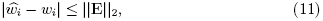

where ||**E**||_2_ is the largest singular value of **E**. To simplify the analysis further, assume **E** = α**Z**, where *α* describes the proportional error in **Z**, so that ||**E**||_2_ = 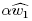. Note that a value of *α* = 0.01 corresponds to a 1% estimation error in **Z**.

In their critique, YC apply the relatedness threshold suggested in GCTA to the Framingham dataset (analyzed in our paper) and report that they do not find the large skews in the singular values we reported in SK. For a threshold of 0.05, although they find the largest singular value (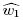 ≈ 500) is of about the same order of magnitude as that reported in our paper, they find that the smallest singular value (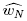 ≈ 110) is substantially larger (we thank Jian Yang for providing us with the numbers).

Suppose that the sampling error in **Z** is small, say 5% (an optimistic value, as we show) i.e., *α* = 0.05 and ||**E**||_2_ = 25. With ||**E**||_2_ = 25 and 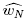 = 110, (11) states that 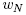 must lie between 85 and 135. But we know that when there is no cryptic relatedness, the smallest singular value of the true GRM is 0 (Part 2 of Theorem 3 in [12]) and therefore, the sampling error must be larger than 5%.

So set *α* = 0.2, for which (11) states that 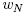 must lie between 10 and 210, but even this interval does not contain 0, and is therefore not large enough. This demonstration shows that the GRM used in GCTA is making estimation errors of at least 20%. These large errors suggest that GCTA’s relatedness threshold cannot,in fact, pick out unrelated individuals.

### 4.2 Trends in Figure 1 of YC

Insights into the singular value trends in figure 1 of YC come from the work of Hoffman [7], who analyzed the effects of cryptic relatedness and population stratification on the eigenvalues of the sample GRM (i.e., squares of the entries in Figure 1 of YC). Hoffman’s results are best summarized through Fig. S1 in his paper (replicated as Fig. 5 below with permission). When cryptic relatedness in the sample is small (panel (a) of Fig. 5), we expect the eigenvalues of the sample GRM to fall sharply towards 0. When the cryptic relatedness in the population is high, we expect to see long stretches of ‘large’ non-zero eigenvalues (panel (b) of Fig. 5, Figure 1 of YC). When there is a combination of the two, we expect the fall to zero to be more gradual (panel (c) of Fig. 5, Figure 3 in SK). Thus the YC results suggest that, even after applying the filtering process suggested in [17], a large number of related individuals is retained in the sample and produce confounding effects above and beyond those described in SK.

**Figure 5:**
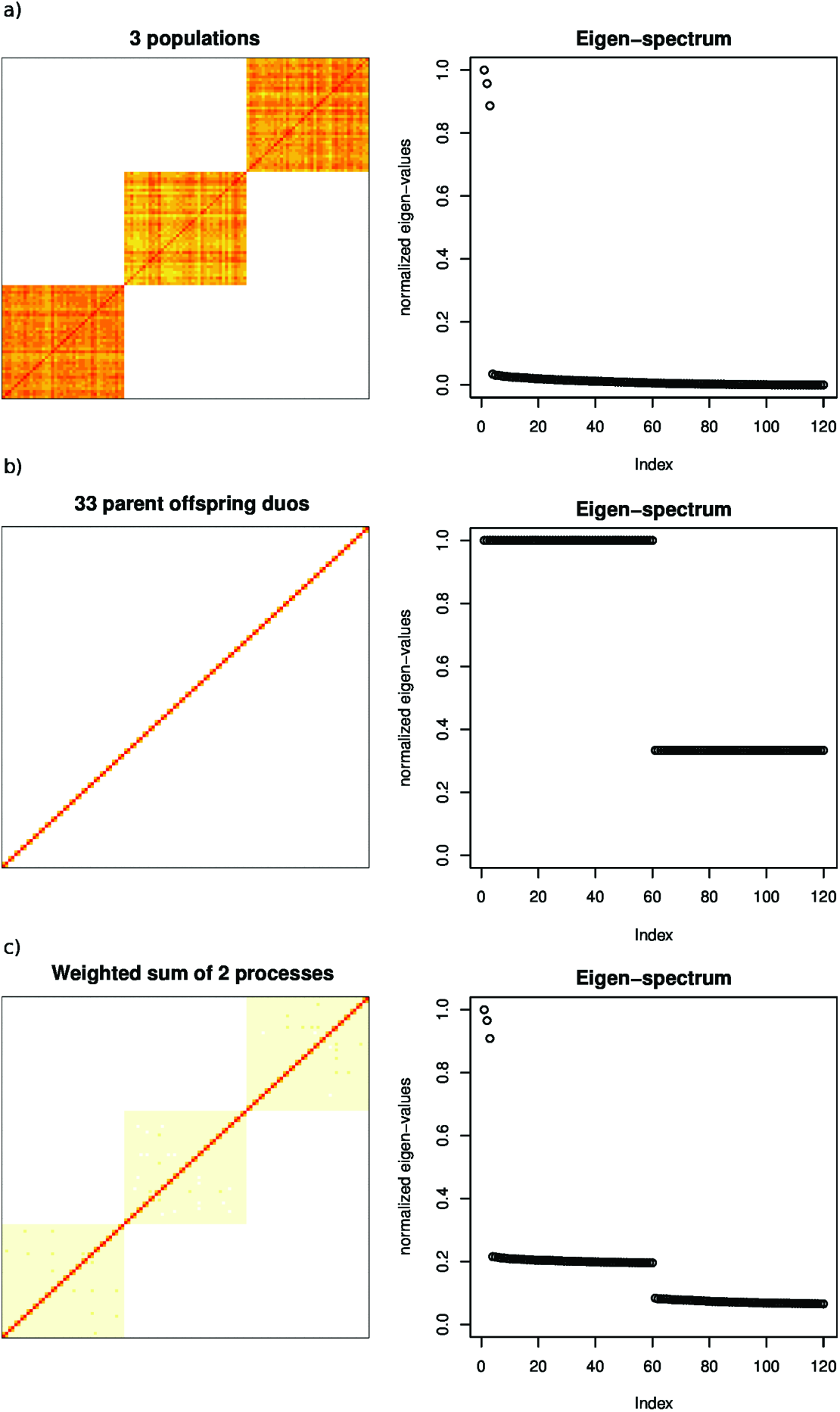
What to expect from the singular(eigen) values of the GRM. Panel (a) shows that when there is stratification and no cryptic relatedness, we expect the eigenvalues to fall to 0 sharply. Panel (b) shows that when the individuals in the population are related, we expect long stretches of non-zero eigenvalues. Panel (c) shows that when there is a combination of the two, we expect the eigenvalues to fall to 0 somewhat gradually. This is supplementary figure S1 in [7]

In Hoffman’s analysis of the Framingham data set (using 3063 individuals), he finds a long stretch of intermediate singular values which he attributes to cryptic relatedness (see section titled ‘Data Analysis’ in [7]). In SK, we did a preliminary filtering using (limited, but reliable) pedigree information to select 2968 ‘unrelated’ individuals for our analysis. As a result, the relatedness in our sample is smaller than that observed in Hoffmann’s; accordingly, the singular values in our plots fall to 0 more rapidly than shown here in Fig. 1. There is clearly some cryptic relatedness in our sample, since the singular values fall to 0 somewhat gradually (similar to panel (c) of Fig. 5). When the cryptic relatedness in the sample is reduced, we expect the skew in the singular values to be larger than that reported in our paper (see the skew for the European population in Fig. 1). In SK we gave GCTA the benefit of the doubt and assumed that the ‘unrelated’ individuals used in the analysis were chosen using more reliable methods than those described in [17]. To demonstrate that the problems with the relatedness threshold are not specific to FHS, we next demonstrate similar problems with the HRS dataset.

### 4.3 The same problem in HRS data

Here we use the HRS dataset to compare the efficiency of kinship coefficients (known to reliably filter unrelated individuals) with the relatedness threshold suggested in [17]s. We show that the latter method, relatedness thresholds, produces much larger errors than a mthod that uses kinship coefficients. Therefore, studies using the relatedness threshold to chose individuals will induce errors beyond those described in SK.

Plots of the singular values of **Z** when subjects are chosen on the basis of kinship coefficients (marked with red dots) and the relatedness threshold (marked with blue dots) are plotted in Fig. 2. Note in Fig. 2 that there is a gap in the singular value spectra obtained using the two methods. Hoffman [7] discusses this gap in his paper and shows that it is to be expected as a consequence of a high-dimensional process like cryptic relatedness.

When the individuals retained in the study are chosen using Kinship Coefficients (see http://hrsonline.isr.umich.edu/sitedocs/genetics/HRS_QC_REPORT_MAR2012.pdf for details), the singular values of **Z** are extremely skewed. In Fig. 2, the largest singular value is 3250, and the smallest singular value is 8 x 10-^6^. For *α* = 10^−4^ (or ||**E**||_2_ = 3250 x 10^−4^ = 0.325), equation (11) states that 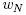 will approximately lie between −0.325 and 0.325 and therefore, the error in estimation of **Z** will be around 0.01%.

On the other hand, when we use a relatedness threshold of 0.025 as suggested in [17] (see Fig. 2 for the scree plot, and Fig. 6 for a histogram of the off-diagonal entries in the GRM), the largest singular value of **Z** is 13216, and the smallest singular value of **Z** is 802. Here, for *α* = 0.05 (or ||**E**||_2_ = 0.05 x 13216 ≈ 660) (11) states that 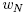 will approximately lie between 802-660 = 142 and 802+660 = 1462. Since this interval does not include 0 and is therefore not large enough. Therefore, the estimation error in **Z** must be larger than 5%.

**Figure 6:**
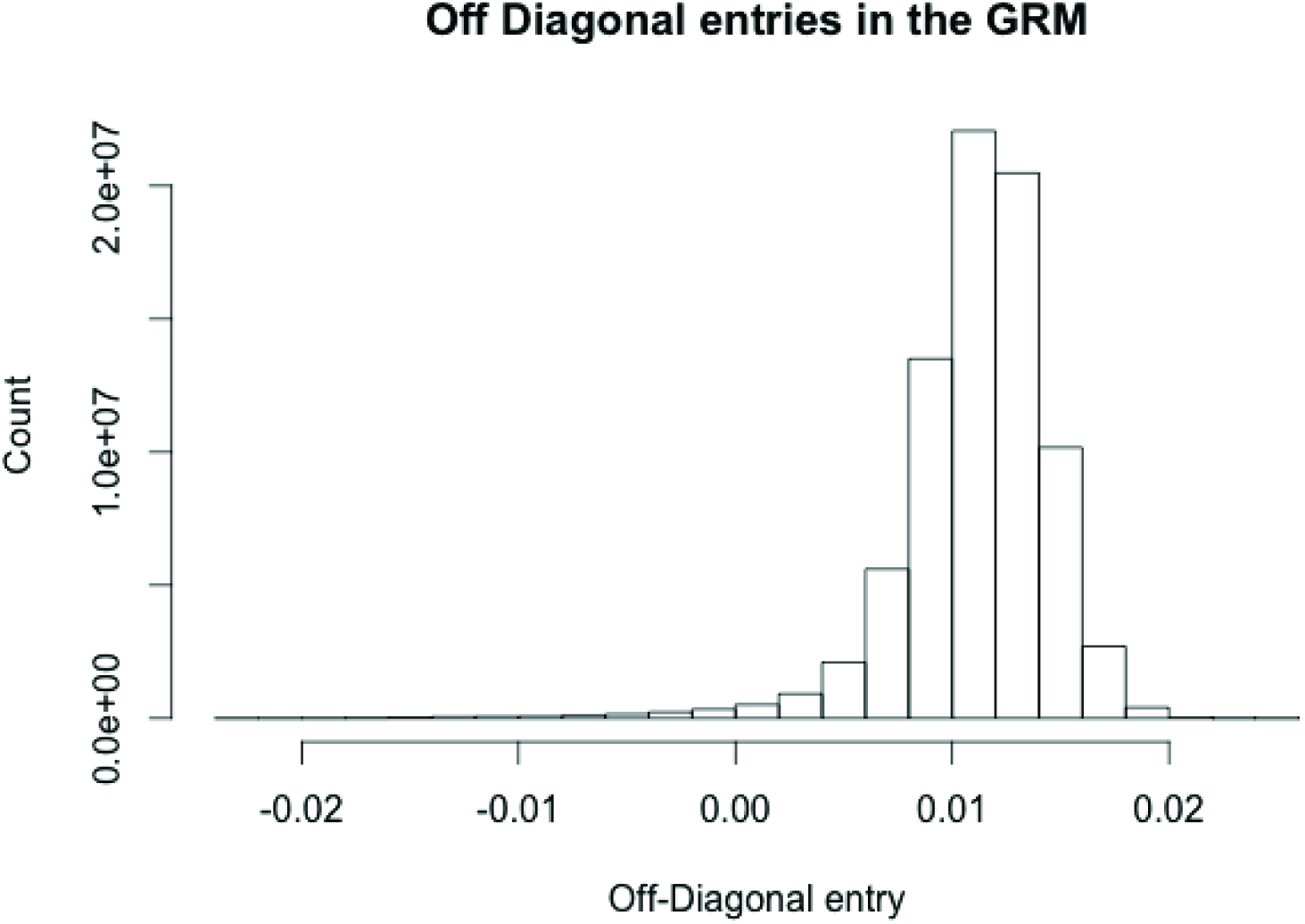
Off-Diagonal entries in the GRM for the HRS dataset after 0.025 thresholding

This illustration shows that the relatedness threshold is an unreliable method for filtering unrelated individuals. In the next section we describe the extreme precision with which **Z** has to be estimated in order for GCTA to work as stated; we will show that even seemingly accurate solutions (e.g., when *α* = 10^−4^) produce extremely inaccurate estimates of heritability.

## 5 Problems with the skew

GCTA uses the singular values 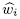 of the sample matrix **Z**, while in reality it should be using the singular values *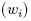* of the ‘true’ matrix **Z**_1_. Our analysis shows that the MLEs depend on terms of the form log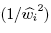. Performing Taylor expansions about the ‘true’ singular value 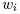, the first order correction in these terms will be of the form

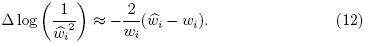

Therefore in essence, the ratio of the error in the singular value to the magnitude of the singular value should be small.

Using (3) and standard results in perturbation theory (see equation 3 in [15]), we can express 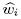 in terms of 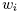 as

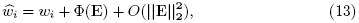

where Φ(**E**) is a linear functional in **E**. Equation (13) shows that a ‘small’ error means that 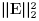 is small; i.e., the square of the largest singular value
of **E** is small, which is consistent with Weyl’s inequality in (11) above. We claimed in SK that the errors associated with the small singular values will be large. To see why, we set **E** = *α**Z*** as in the previous section. For this setting, 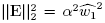. From (13), for the errors in the smallest singular value 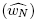 to be small, we require

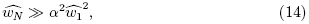

or

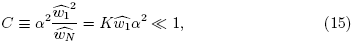

where *K = 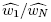* is the condition number of the matrix.

Inserting 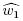 = 1000, 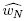 = 100, and *α* = 0.01 in (15) gives *C* = 1; this shows that even when the smallest singular value is ‘large’ and the condition number is modest, extreme precision is required for proper estimation of **Z**_1_. Further, this example demonstrates that the sampling error in the smaller singular values will be large so long as they are small *relative* to the largest singular values; the absolute magnitude has little relevance.

For the HRS dataset, with *α* = *O*(10^−4^), 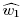 = *O*(10^3^), and 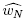 = *O*(10^−6^), C = *O*(10^4^); therefore, a sampling error of 0.01% in **Z** is extremely large. This illustration demonstrates the very (unreasonably?) high accuracy required in **Z** for the GREML to work as described in YC.

When the singular values of **Z** are skewed, the precision required in the GRM, **A**, is much higher than the precision required in **Z**. To see why, we analyze the conditions under which the errors in the eigenvalues of the GRM 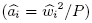 will be small. From (14), for the errors in the smallest eigenvalue of the GRM to be small, we require the sampling error in the GRM described in (4) (**F** = *β***A**) to approximately satisfy

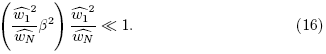

Comparing (15) and (16) shows that

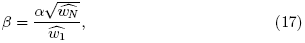

which means that a high precision in **Z** corresponds to a much higher precision in **A**.

In the context of the HRS dataset described above, *α* = 10^−4^ (which we showed was too large) corresponds to *β* = *O*(10^−8^). This implies that an error of 10^−6^% in the GRM would be too large.

It has been claimed in applying GCTA, that the errors they find in the GRM are small; the problem is that they are not small enough. For instance in supplementary figure S1 of [16], the diagonal entries in the sample GRM whose ‘true’ value is 1 are plotted and entries as large as 1.1 are found, correspond to an error of 10%. This plot clearly shows that contrary to the claims in YC, the sampling error in the GRM are not small, and problems with the sampling errors are bound to exist.

## 6 Acknowledgements

SKK is funded by the Stanford Center for Computational, Evolutionary and Human Genomics. D.H.R. is supported by National Institute on Aging Grant K01AG047280. This research uses data from the HRS, which is sponsored by the National Institute on Aging (Grants NIA U01AG009740, RC2AG036495, and RC4AG039029) and conducted by the University of Michigan.

## References

[1] P. J. Bickel and E. Levina. Covariance regularization by thresholding. The Annals of Statistics, pages Vol.36, No. 6 2577–2604, 2008.

[2] P. J. Bickel and E. Levina. Regularized estimation of large covariance matrices. The Annals of Statistics, pages Vol. 36, No.1 199–227, 2008.

[3] T. T. Cai, Z. Ren, and H. H. Zhou. Estimating structured high-dimensional covariance and precision matrices: Optimal rates and adaptive estimation. The Annals of Statistics, 38:2118–2144, 2014.

[4] Cross-Disorder Group of the Psychiatric Genomics Consortium Genetic relationship between five psychiatric disorders estimated from genome-wide snps. Nature genetics, 45(9):984–994, 2013.

[5] T. Hastie, R. Tibshirani, and J. Friedman. The elements of statistical learning, volume 2. Springer, New York, 2009.

[6] C. R. Henderson. Best linear unbiased estimation and prediction under a selection model. Biometrics, pages 423–447, 1975.

[7] G. E. Hoffman. Correcting for population structure and kinship using the linear mixed model: theory and extensions. PloS One, 8(10):e75707, 2013.

[8] A. Izenman. Modern multivariate statistical techniques, volume 1. Springer, New York, 2008.

[9] S. K. Kumar, M. W. Feldman, D. H. Rehkopf, and S. Tuljapurkar. Limitations of gcta as a solution to the missing heritability problem. Proceedings of the National Academy of Sciences U S A, 113(1):E61–E70, 2016.

[10] O. Ledoit and M. Wolf. Honey, I shrunk the sample covariance matrix. J Portf Manag, 30(4), 2004.

[11] S. H. Lee, N. R. Wray, M. E. Goddard, and P. M. Visscher. Estimating missing heritability for disease from genome-wide association studies. The American Journal of Human Genetics, 88(3):294–305, 2011.

[12] N. Patterson, A. L. Price, and D. Reich. Population structure and eigenanalysis. PLoS genetics, 2(12):e190, 2006.

[13] R. L. Quaas and E. Pollak. Mixed model methodology for farm and ranch beef cattle testing programs. Journal of Animal Science, 51(6):1277–1287, 1980.

[14] G. K. Robinson. That BLUP is a good thing: the estimation of random effects. Statistical science, 6, 1991.

[15] G. W. Stewart. Perturbation theory for the singular value decomposition. Institute for Advanced Computer Studies, University of Maryland, College Park, MD., 1998.

[16] A. A. Vinkhuyzen, N. L. Pedersen, J. Yang, S. H. Lee, P. K. Magnusson, W. G. Iacono, M. McGue, P. Madden, A. C. Heath, M. Luciano, et al. Common snps explain some of the variation in the personality dimensions of neuroticism and extraversion. Translational psychiatry, 2(4):e102, 2012.

[17] J. Yang, B. Benyamin, B. P. McEvoy, S. Gordon, A. K. Henders, D. R. Nyholt, P. A. Madden, A. C. Heath, N. G. Martin, G. W. Montgomery, et al. Common SNPs explain a large proportion of the heritability for human height. Nature genetics, 42(7):565–569, 2010.

[18] J. Yang, S. H. Lee, N. R. Wray, M. E. Goddard, and P. M. Visscher. Commentary on” limitations of gcta as a solution to the missing heritability problem”. bioRxiv, page 036574, 2016.

[19] J. Yang, T. A. Manolio, L. R. Pasquale, E. Boerwinkle, N. Caporaso, J. M. Cunningham, M. de Andrade, B. Feenstra, E. Feingold, M. G. Hayes, et al. Genome partitioning of genetic variation for complex traits using common snps. Nature genetics, 43(6):519–525, 2011.

